# FLUAV RAM-IGIP: A modified live influenza virus vaccine that enhances humoral and mucosal responses against influenza

**DOI:** 10.1101/2024.01.23.576908

**Authors:** C. Joaquín Cáceres, L. Claire Gay, Aarti Jain, Teresa D. Mejías, Matias Cardenas, Brittany Seibert, Flavio Cargnin Faccin, Brianna Cowan, Ginger Geiger, Amy Vincent Baker, Silvia Carnaccini, D. Huw Davies, Daniela S. Rajao, Daniel R. Perez

## Abstract

Current influenza A vaccines fall short, leaving both humans and animals vulnerable. To address this issue, we have developed attenuated modified live virus (MLV) vaccines against influenza using genome rearrangement techniques targeting the internal gene segments of FLUAV. The rearranged M2 (RAM) strategy involves cloning the M2 ORF downstream of the PB1 ORF in segment 2 and incorporating multiple early stop codons within the M2 ORF in segment 7. Additionally, the IgA-inducing protein (IGIP) coding region was inserted into the HA segment to further attenuate the virus and enhance protective mucosal responses. RAM-IGIP viruses exhibit similar growth rates to wild type (WT) viruses in vitro and remain stable during multiple passages in cells and embryonated eggs. The safety, immunogenicity, and protective efficacy of the RAM-IGIP MLV vaccine against the prototypical 2009 pandemic H1N1 strain A/California/04/2009 (H1N1) (Ca/04) were evaluated in Balb/c mice and compared to a prototypic cold-adapted live attenuated virus vaccine. The results demonstrate that the RAM-IGIP virus exhibits attenuated virulence in vivo. Mice vaccinated with RAM-IGIP and subsequently challenged with an aggressive lethal dose of the Ca/04 strain exhibited complete protection. Analysis of the humoral immune response revealed that the inclusion of IGIP enhanced the production of neutralizing antibodies and augmented the antibody-dependent cellular cytotoxicity response. Similarly, the RAM-IGIP potentiated the mucosal immune response against various FLUAV subtypes. Moreover, increased antibodies against NP and NA responses were observed. These findings support the development of MLVs utilizing genome rearrangement strategies in conjunction with the incorporation of immunomodulators.

**IMPORTANCE:** Current influenza vaccines offer suboptimal protection, leaving both humans and animals vulnerable. Our novel attenuated MLV vaccine, built by rearranging FLUAV genome segments and incorporating the IgA-inducing protein, shows promising results. This RAM-IGIP vaccine exhibits safe attenuation, robust immune responses, and complete protection against lethal viral challenge in mice. Its ability to stimulate broad-spectrum humoral and mucosal immunity against diverse FLUAV subtypes makes it a highly promising candidate for improved influenza vaccines.

## INTRODUCTION

Seasonal influenza epidemics caused by influenza A (FLUAV) and influenza B (FLUBV) viruses are responsible for 3-5 million severe respiratory illnesses and approximately 600,000 deaths worldwide each year (1). In the United States alone, influenza infections result in an average economic burden of $87 billion due to preventive and therapeutic treatments, hospitalization costs, and lost workdays (2). Vaccination serves as the primary defense against influenza infections, but the efficacy of current vaccines has raised concerns, necessitating the development of more effective vaccines. The FDA has approved three types of influenza vaccines for human use: inactivated vaccines (IV), recombinant influenza protein vaccines (RIV), and live attenuated influenza virus vaccines (LAIV). IV and RIV vaccines elicit strong humoral immune responses but provide limited or no cellular immune response activation. LAIV vaccines are a form of modified live virus (MLV) vaccines that carry temperature-sensitive, cold-adapted mutations. The FLUAV components of the currently approved LAIV vaccines for human use include those based on the cold-adapted A/Ann Arbor/6/60 strain and the cold-adapted A/Leningrad/134/17/57 strain (caLen). Like MLVs, LAIVs mimic a natural influenza infection, eliciting a combination of humoral and cellular immune responses, making them theoretically more effective influenza vaccines due to their multidimensional responses (3). Recent issues have limited the full potential of LAIVs. Quadrivalent LAIV formulations introduced since the 2013-2014 influenza season have demonstrated low efficacy in protecting against seasonal influenza viruses, particularly in children (4). Thus, the need to explore alternative MLV strategies against influenza.

FLUAV and FLUBV viruses are amenable to substantial genome rearrangements, with the PB1 segment tolerating the addition of various foreign sequences, specifically M2 (RAM) and NEP (RANS) (5, 6). Modifications in segment 7 to prevent expression of M2 in RAM or in segment 8 to prevent expression of NEP in RANS further attenuate these viruses (5, 6). FLUAV and FLUBV with RAM or RANS rearrangements induced protection against influenza virus challenge in different mammalian models (5, 6). Unlike current LAIVs, the genome rearrangement MLV strategy can be used to update the entire vaccine virus backbone.

Stimulating mucosal immune responses is critical for developing effective MLVs against influenza. Following influenza infection, IgA and IgG responses are detected in the airway mucosa, exhibiting neutralizing activity against influenza. IgA, particularly secretory IgA (sIgA) in its multimeric forms, is typically more broadly neutralizing than IgG (7, 8). IgA neutralizes pathogens without causing inflammation due to its inability to fix and activate the complement cascade (7, 8). Recognizing the advantages of stimulating mucosal responses, we have investigated the role of the IgA-inducing protein (IGIP) (9) as a natural adjuvant incorporated into the genome of recombinant MLVs. Initially characterized in bovine gastrointestinal-associated lymphoid tissue (GALT), IGIP positively regulates IgA expression and is highly conserved among mammals, with a predicted molecular weight between ∼5.1 and ∼5.9 KDa (10, 11). To evaluate the role of IGIP in the context of MLVs, we engineered a chimeric segment 4 that expresses IGIP and HA. We have previously reported on an attenuated FLUAV vaccine carrying IGIP in a virus backbone with temperature sensitive mutations (att-IGIP) that resulted in improved safety profiles while enhancing humoral and mucosal immune responses in mice (12). Studies in pigs vaccinated with a bivalent FLUAV att-IGIP against H3N2 and H1N1 swine FLUAVs revealed protection after virus challenge and complete block in virus transmission in pigs compared to partial transmission in pigs immunized with a bivalent FLUAV att non-IGIP-vaccine(13).

This report further explores the potential of IGIP as a natural adjuvant in the context of the FLUAV RAM MLV strategy using the mouse model. FLUAV RAM-IGIP (H1N1) was generated. Both viruses were stable *in vitro* over serial passages in eggs and/or tissue culture cells and grew to titers like the prototypical WT Ca/04 strain. Mice vaccinated intranasally with the FLUAV RAM-IGIP showed no clinical signs and were completely protected against lethal challenge with Ca/04. Analyses of the humoral and mucosal responses revealed higher production of neutralizing antibodies and mucosal responses in FLUAV RAM-IGIP-vaccinated mice than those vaccinated with the caLen (H1N1) control.

## RESULTS

### In vitro properties of FLUAV RAM and FLUAV RAM-IGIP MLVs

The FLUAV RAM attenuating strategy involves inserting the M2 ion channel ORF downstream of the PB1 ORF (segment 2) and separating them with a spacer encoding the Thosea asigna virus 2A protease (2ATaV), whose activity is predicted to release M2 from PB1 during translation. Multiple early stop codons are introduced in segment 7 to eliminate M2 expression(6). We evaluated the polymerase activity mediated by the RAM PB1-M2 gene derived from the A/California/04/2009 (H1N1) (Ca/04) strain(14) in a minigenome assay at different temperatures (Fig S1) and compared it to the previously described platform with temperature sensitive mutations (att)(15). Polymerase activity from the RAM and att polymerase complexes were lower than the WT polymerase complex, regardless of temperature. However, differences were only statistically different between the WT vs. att control at 39°C (p=0.016).

We generated a FLUAV RAM internal gene segment backbone using modified PB1-M2 and M_esc_ segments derived from Ca/04 and the remaining unmodified internal segment constellation (PB2, PA, NP, and NS) from OH/04, a previously described virus vaccine backbone(15). The H1 HA segment derived from Ca/04 was modified by introducing a 56 amino acid sequence of which 24 amino acids correspond to the matured IGIP peptide sequence(12). IGIP was cloned downstream of the HA’s signal peptide sequence but upstream of the mature HA ORF. Additional amino acid sequences were included downstream of IGIP to encode proteolytic activities that release IGIP from HA during translation(12). The resulting virus, RAM-IGIP, and its non-IGIP counterpart RAM were evaluated for replication fitness *in vitro* at 35°C in MDCK cells and in differentiated human-airway epithelial (HAE) cells (16). The WT Ca/04 and att (H1N1)(15) viruses were included as comparators. The RAM and RAM-IGIP viruses replicated well in MDCKs (Fig 1A). Statistically significant differences versus the WT Ca04 were observed at the 12, 24, and 48 hpi marks (p<0.05) but not at the 72 hpi time point. Of note, the IGIP modification in HA did not significantly affect the growth of the RAM backbone compared to non-IGIP counterpart. In HAE cells (Fig 1B), there were no statistically significant differences observed among the different viruses evaluated. Additional studies in MDCK cells showed lower virus yields at 35, 37, 39, and 41°C compared to the WT Ca/04 virus. However, only at 39 and 41°C were these differences statistically significant, although the effect was not as profound as with the att (H1N1) virus (Fig 1C)

**Fig 1.**
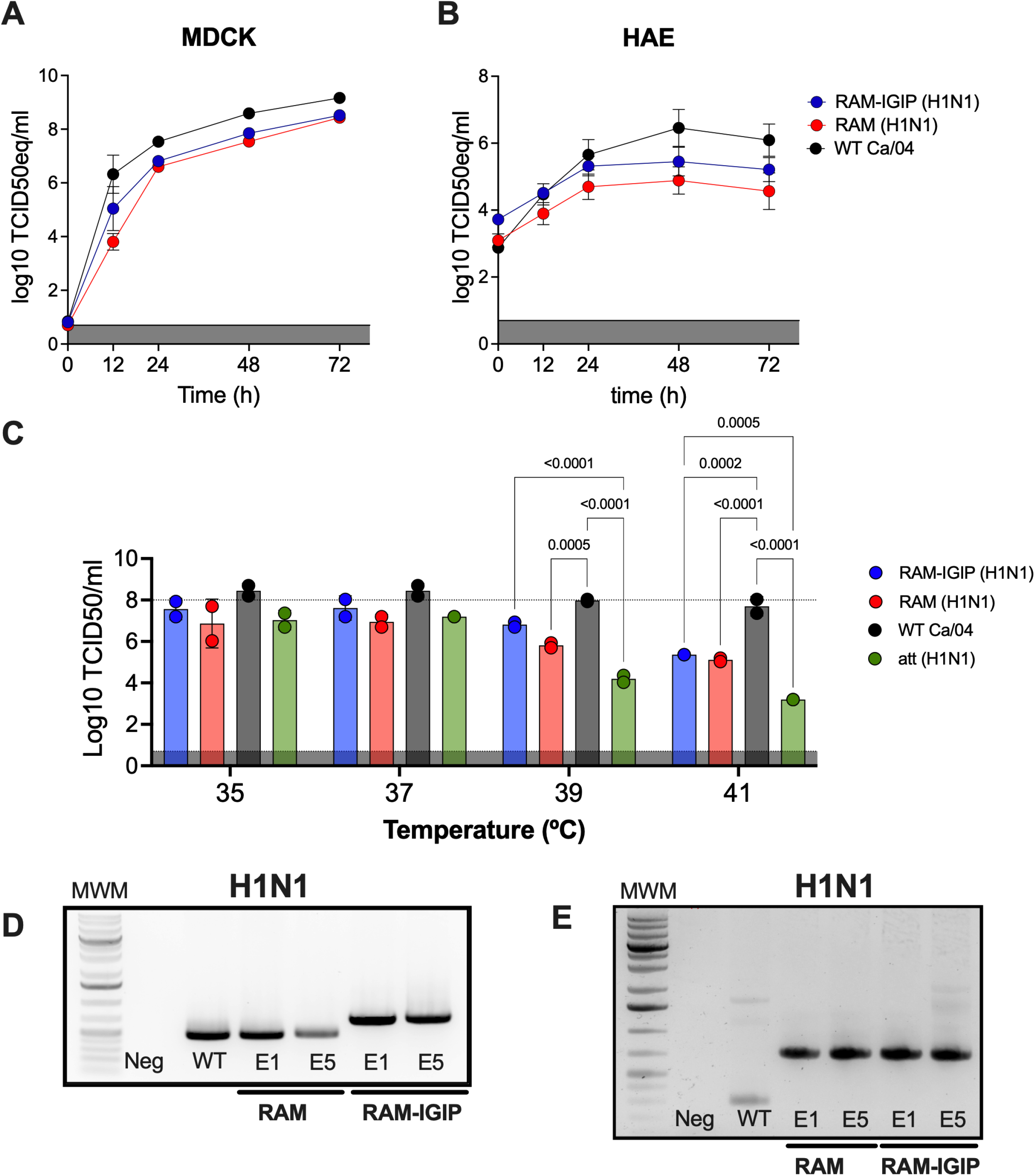
In vitro properties of MLV vaccines. *In vitro,* growth kinetics were performed with RAM-IGIP, RAM, WT H1N1 virus in **(A)** MDCK and **(B)** human airway epithelial cells (HAE). Stability analysis of the different RAM viruses after serial passages in eggs. **(C)** Growth profiles at different temperatures of RAM-IGIP and RAM viruses compared to WT Ca/04 and a temperature sensitive Ca/04 virus (att) inoculated in MDCKs. Titers were analyzed at 72 hpi, established by TCID50, and expressed as Log10 TCID50/mL. The stability of **(D)** HA and **(E)** PB1-M2 in RAM-IGIP-H1N1 viruses was established after 5 serial passages of viruses in eggs (E5) and determined by RT-PCR analyses. Controls include WT virus and mock-inoculated allantoic fluid and virus from the E1 passage.

### FLUAV RAM and FLUAV RAM-IGIP MLVs are stable during serial passage

Passage of the RAM MLVs in eggs five times demonstrated the stability of all modified components in the genome (Fig 1D-E), as confirmed by PCR analysis. The estimated amplicon size for IGIP-H1 is 699 bp and for the H1 WT is 477 bp. The expected amplicon sizes for PB1-M2 is 679 bp and for PB1 WT is 273 bp. This observation aligns with a similar strategy for FLUBV(6). Consistent with a previous report (12), the inclusion of IGIP in the HA remained stable (Fig 1D). Sequence analysis corroborated the PCR findings for the various modified segments. Additionally, sequence analysis of segment 7 revealed a single nucleotide change at position 863 (t863c) in the M segment downstream of the M1 ORF and the introduced early stop codons. This change is located downstream of the M1 ORF, and its location relative to the early stop codons suggests that its phenotypic contribution is negligible.

### FLUAV RAM and FLUAV RAM-IGIP MLVs are attenuated in mice

To evaluate the safety profiles of the RAM-IGIP and RAM strains, Balb/c mice (n=24/group, ½ female) were inoculated with 10^5^ TCID50 of virus/mouse. Control groups included mice inoculated with a cold adapted LAIV (H1N1) vaccine based on the A/Leningrad/134/17/57 strain (caLen, n=24/group, ½ female) (17), mice inoculated with WT Ca/04 (n=10, ½ female), and mice inoculated with PBS (mock, n=12, ½ female). Assessment of clinical signs (Fig 2A) revealed no changes in activity or physical appearance in mice inoculated with the RAM-IGIP, RAM, and caLen viruses compared to the mock group. Consistent with these observations, mice inoculated with the modified viruses exhibited no significant body weight changes and 100% survival (Fig 2B). In contrast, mice inoculated with WT Ca/04 began showing clinical signs at 2 days post-inoculation (dpi), including decreased activity, reduced responsiveness to stimulation, rough fur, body weight loss (≥25% by 5 dpi) and were humanely euthanized.

**Fig 2.**
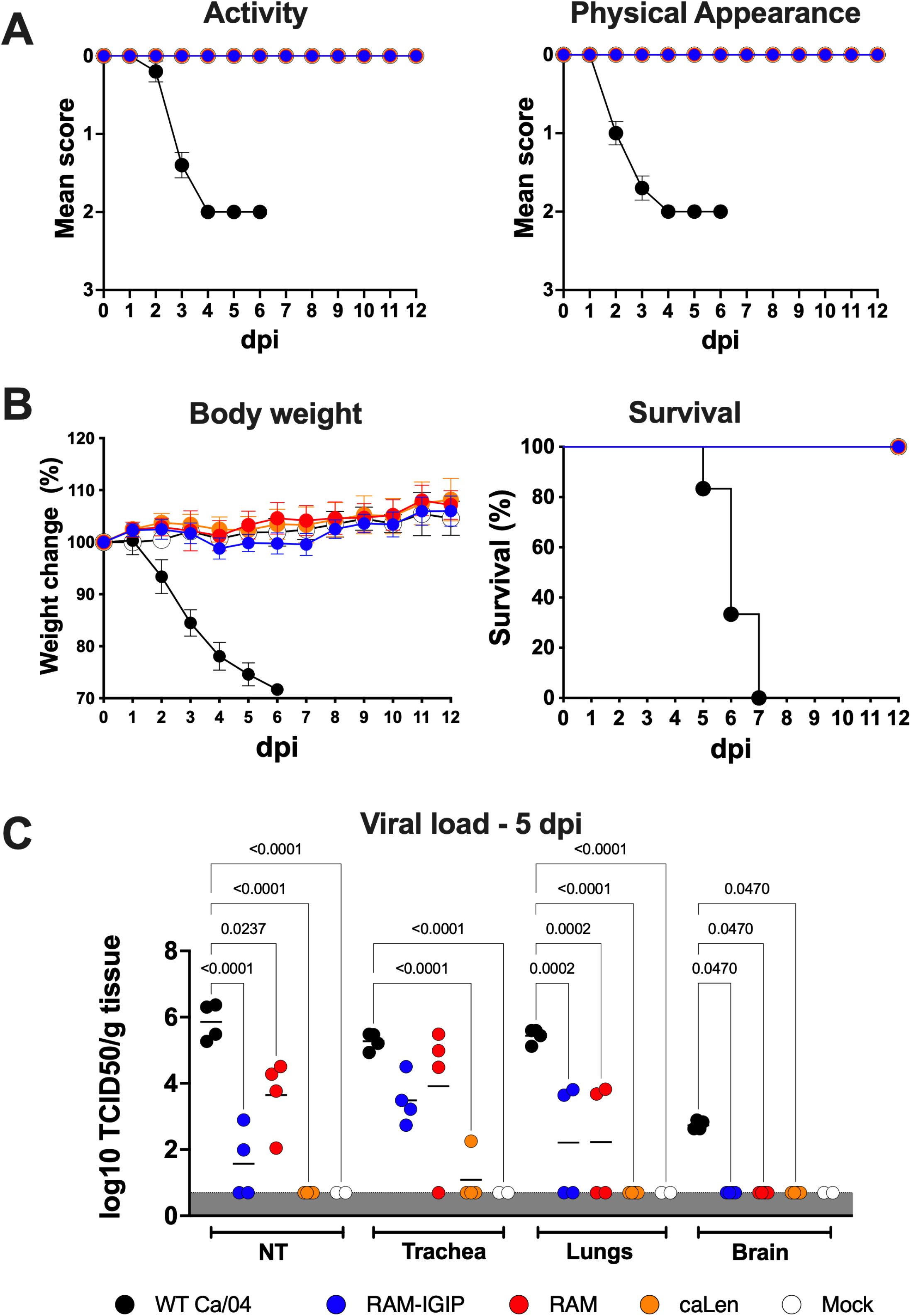
Attenuation of the RAM viruses *in vivo*. Mice were inoculated I.N. with 10^5^ TCID50/mouse of WT Ca/04 (black), RAM-IGIP (blue), RAM (red), caLen (orange) or mock-inoculated (white) **(A)** Activity and Physical Appearance **(B)** Body weight and Survival were monitored for 12 days after vaccination. **(C)** A subset of animals was sacrificed from each group at 5 dpi and live virus titers in nasal turbinates (NT), trachea, lungs, and brain were quantified and expressed as Log_10_ TCID50/gr of tissue homogenate.

Viral loads and innate immune responses in samples collected at 5 dpi were analyzed (n=4/group, n=2/mock group, ½ female). Compared with WT-inoculated mice, significant reductions in viral loads were detected in nasal turbinates (NT), trachea, lungs, and brain in the RAM-IGIP and RAM groups (Fig 2C). Innate immune responses were analyzed using a 26-plex Luminex assay on samples from lungs and serum. Out of the analyzed cytokines, only a few exhibited statistically significant differences between vaccinated groups and the WT Ca/04 virus control. Specifically, MCP-1, IL-18, IL-12 (p70), and TNF-α serum levels in vaccinated groups were comparable to the mock control group and significantly lower than those in the WT Ca/04 group (Fig 3). RAM and caLen groups displayed statistically lower GM-CSF serum levels compared to the WT Ca/04 group, with a similar trend in the RAM-IGIP group (Fig 3). All vaccinated groups showed lower serum levels of IL-6 and MCP-3 compared to the WT Ca/04 group, with only caLen reaching statistical significance. The remaining cytokines showed trends dependent on the vaccine group but were not statistically significant (not shown). These observations are consistent with the attenuated phenotype of the vaccine viruses (Fig 2). Histopathological analysis (Fig S2 A-B) of the groups at 5 dpi revealed severe suppurative tracheitis and bronchopneumonia with accumulation of abundant intraluminal exudate in all mice inoculated with WT Ca/04. These also presented mild, sparse infiltration of lymphocytes in the peri bronchial lamina propria and adjacent interstitium. These lesions corresponded to high positivity of virus antigen likely matching the extent of virus replication in these tissues. In contrast, the RAM-IGIP and RAM groups had predominantly a non-suppurative, mild to moderate, lymphoplasmacytic infiltration in the tracheal and pulmonary bronchial mucosa.

**Fig 3.**
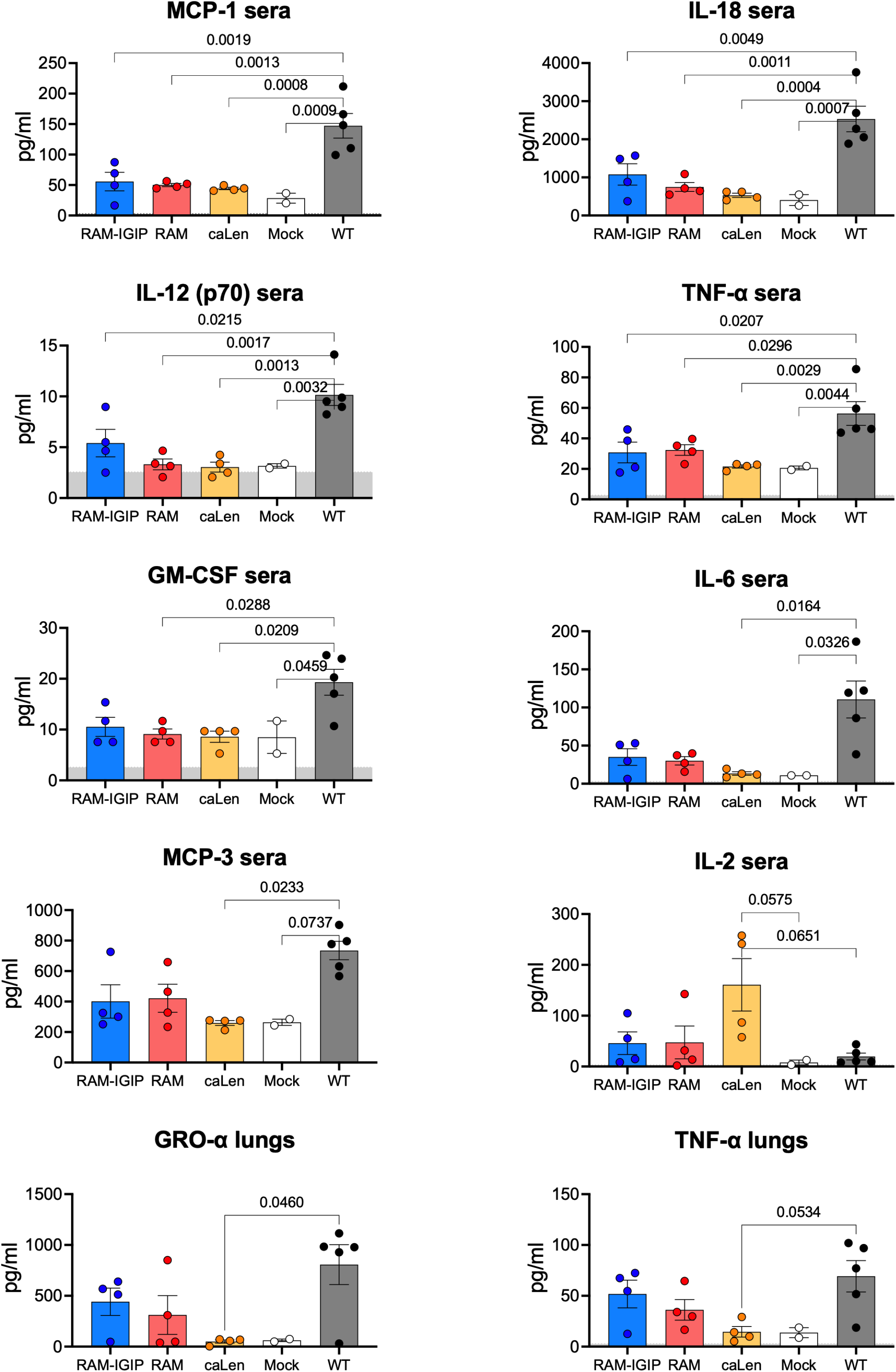
Innate immune responses post-infection. Serum and lung cytokines were detected using a 26-plex Luminex assay in samples collected at 5 dpi. Only those with statistically significant differences among groups are shown. Values in y-axis correspond to pg/mL.

Inflammation was associated with minimal necrosis of the tracheal and bronchial respiratory epithelium and rarely pneumocytes type I. This corresponded to the presence of small amounts of virus antigens. This is expected as live vaccines rely on virus replication to develop memory and cell-mediated immune response. Lesions were absent in the tissues of the caLen group, and virus antigens were too low to detect at immunohistochemistry.

### FLUAV RAM and FLUAV RAM-IGIP MLVs confer protection against lethal homologous challenge in Balb/c mice

Three weeks after a single dose MLV or caLen inoculation, a subset of mice (n=12/group, ½ female) were challenged with 1,000 times the lethal dose 50 of WT Ca/04 (Fig 4) (12, 14). Controls included mock-vaccinated-challenged mice (WT Ca/04, n=12/group, ½ female) and mock-vaccinated-non-challenged mice (mock, n=6/group, ½ female). Following challenge, mice previously vaccinated with either the RAM-IGIP or RAM strains exhibited normal activity and physical appearance, absence of clinical signs, no body weight loss, and 100% survival (Fig 4A and B). Mice previously inoculated with caLen maintained normal physical appearance but experienced mild reductions in activity and significant body weight changes between days 3 and 5 post-challenge (dpc, day 3; p=0.015, day 4; p=0.0005, and day 5; p<0.0001, Fig 4B). However, all caLen-vaccinated mice survived. As expected, starting at 2 dpc, mock-vaccinated-challenged mice (WT Ca/04) exhibited a rapid decline in activity, deterioration in physical appearance, and significant body weight loss, with no survivors by 7 dpc.

**Fig 4.**
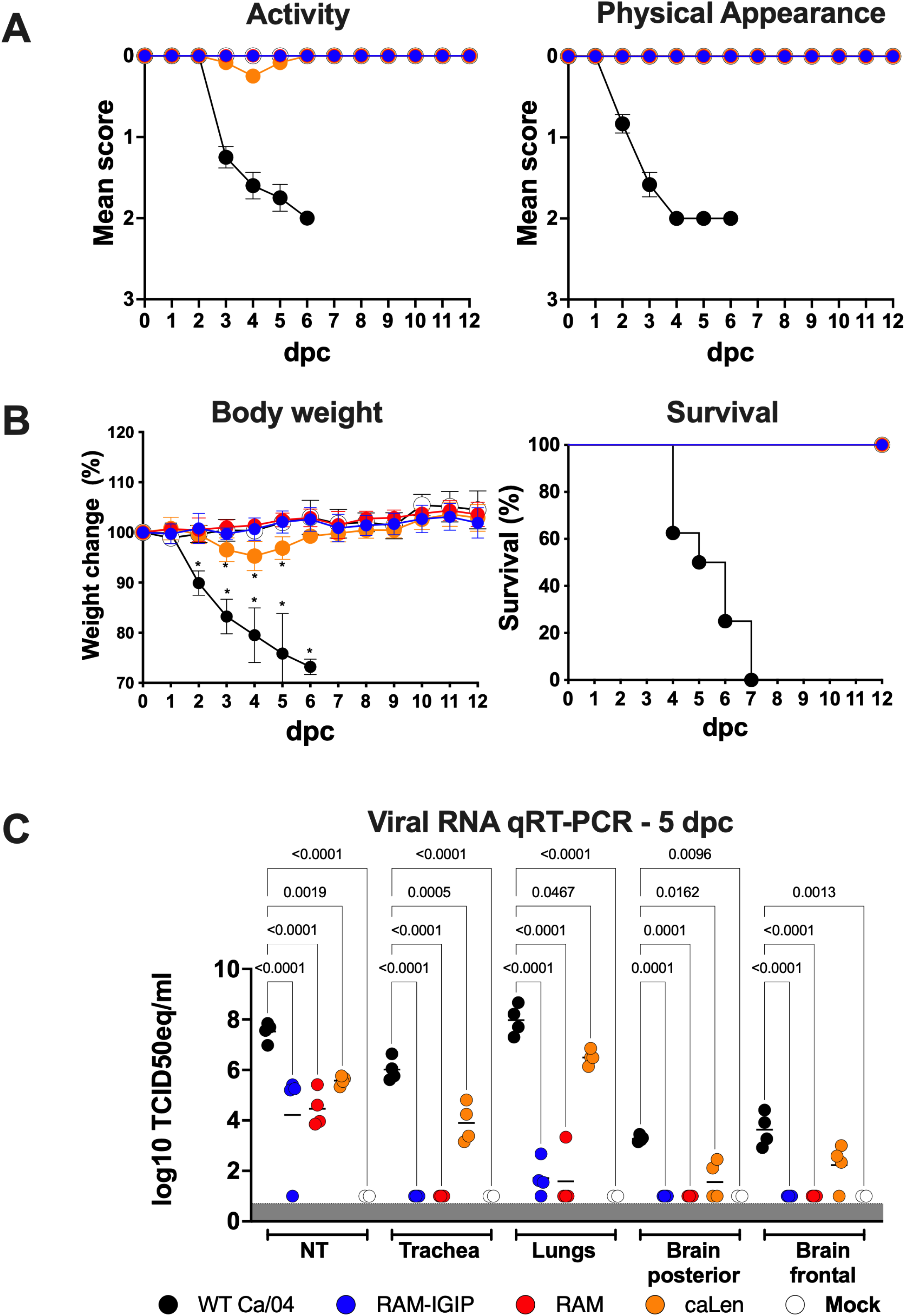
Efficacy of vaccines against homologous lethal challenge. Mice were vaccinated I.N. with RAM-IGIP (blue), RAM (red), caLen (orange) or mock-vaccinated. At three weeks post-vaccnation mice were were challenged with 1000 MLD50 of Ca/04(14). Controls included mock vaccinated-challenged (WT Ca/04, black) and mock vaccinated-non challenged (mock, white) controls. **(A)** Activity and physical appearance and **(B)** body weight and survival were monitored for 12 days after the challenge (dpc).A subset of mice was sacrificed at 5 dpc and vRNA loads in nasal turbinates (NT), trachea, lungs, and brain were quantified and expressed as Log_10_ TCID50 equivalents/mL (log_10_ TCID50 eq/mL).

Viral vRNA loads were evaluated using RT-qPCR at 5 dpc in a subset of mice from each group following challenge (Fig 4C). Overall, challenge virus vRNA levels were significantly lower in either RAM-IGIP- or RAM-vaccinated group compared to mock vaccinated-challenged mice. After challenge, higher vRNA loads were observed in caLen-vaccinated mice compared to either RAM- or RAM-IGIP-vaccinated groups. Despite vRNA detection, mice in the RAM-IGIP, RAM, and caLen groups showed undetectable live virus in nasal turbinates (NT), trachea, and lungs, whereas mock vaccinated-challenged mice (WT Ca/04) exhibited high viral loads, with titers between 10^5^ and 10^7^ TCID50/gr of tissue (not shown).

At 5 days following virus challenge, cytokine levels in lung homogenates (Fig. 5) from mice in the RAM-IGIP and RAM groups were significantly lower than those in the caLen and mock vaccinated-challenged (WT Ca/04) groups. Notably, these levels in the RAM-IGIP and RAM groups were comparable to those observed in the mock-vaccinated-non-challenged control group, indicating a potential dampening of the inflammatory response (Fig. 5). Cytokines exhibiting significantly lower levels in the RAM-IGIP and RAM groups included: IL-1β, IP-10, MIP-1α, GM-CSF, IL-17A, TNF-α, MIP-1β, MCP-3, IL-22, GRO-α, and MCP-1. Furthermore, levels of IL-6, IL-18, IFN-ɣ, IL-5, and IL-12 (p70) were significantly lower in all vaccinated groups compared to the mock-vaccinated-challenged WT Ca/04 group. Conversely, IL-23, IL-10, and IL-2 levels were significantly lower only in samples from the RAM-IGIP group compared to the WT Ca/04 group. Lung homogenates exhibited additional cytokines with diverse expression patterns (Fig S3). Post-challenge, IL-5 levels were significantly lower while RANTES levels were significantly higher in all vaccinated groups compared to WT Ca/04 controls (Fig S3). Additionally, eotaxin, and MIP-2 levels were elevated in the caLen group compared to non-challenged mock controls, reaching levels like those observed in the WT Ca/04 group. In serum samples (Fig 6) and compared to vaccinated mice, the mock-vaccinated-challenged WT Ca/04 group exhibited significantly higher levels of IL-18, MCP-3, MCP-1, IL-12(p70), and IL-6. Compared to the mock-vaccinated-challenged WT Ca/04 group, IP-10 and IFN-ɣ were significantly lower in the RAM-IGIP and RAM groups, but not in the caLen group (Fig 6), whereas TNF-α and GM-CSF were lower in the RAM and caLen groups but not in the RAM-IGIP group. IL-22 was marginally significantly lower only in the RAM group compared to the WT Ca/04 group. Other serum cytokines trend lower in vaccine groups compared to the WT Ca/04 group, but differences were not statistically significant (not shown, data available upon request). Overall, the lower pro-inflammatory cytokine levels in either RAM-IGIP or RAM groups compared to either caLen or mock groups were consistent with the significantly lower vRNA detection (Fig 4). The histopathological analysis at 5 dpc (Fig S2 C-D) revealed severe suppurative tracheitis, bronchopneumonia, and abundant virus antigen detection in the mock-vaccinated-challenged WT Ca/04 group. In the RAM-IGIP and RAM groups, lymphoplasmacytic response predominated in lungs and tracheas, while necrotizing and suppurative lesions were minimal. Lymphocytes and plasma cells were organized in dense sheets, or clustering to form lymphoid nodules in the mucosal lamina propria of the main airways or collaring vessels. These may be signs of an organized cell-mediated immune response, likely prompted by vaccination and then triggered by the challenge. Lesions were far more severe in the caLen group than in any other vaccine groups, resembling the WT Ca/04 group. The lymphoplasmacytic response was present to a greater extent than in WT Ca/04, as expected from a vaccination group; however, lymphocytes were mostly dispersed in the tissue and infiltrating affected respiratory epithelium or collaring vessels. These findings corresponded to the presence of lower amounts of virus antigens than in the WT Ca/04. No virus antigens were detected in either the RAM-IGIP or the RAM groups by immunohistochemistry.

**Fig 5.**
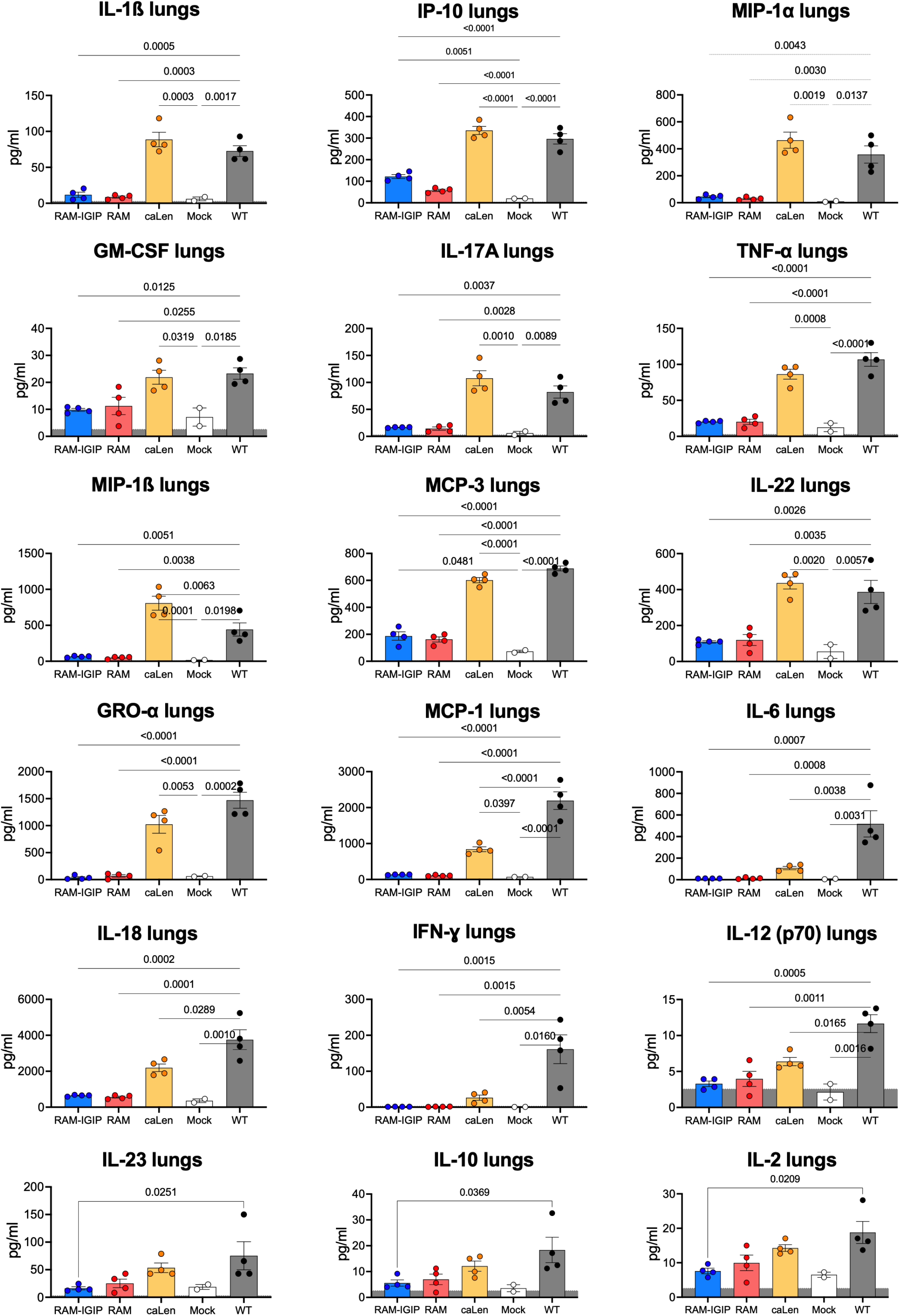
Innate immune responses in lungs after virus challenge. Lung cytokine levels were analyzed in samples collected at 5 dpc. Only those with statistically significant differences among groups are shown. Values in y-axis correspond to pg/mL. Statistically significant differences are shown with p values above lines or brackets.

**Fig 6.**
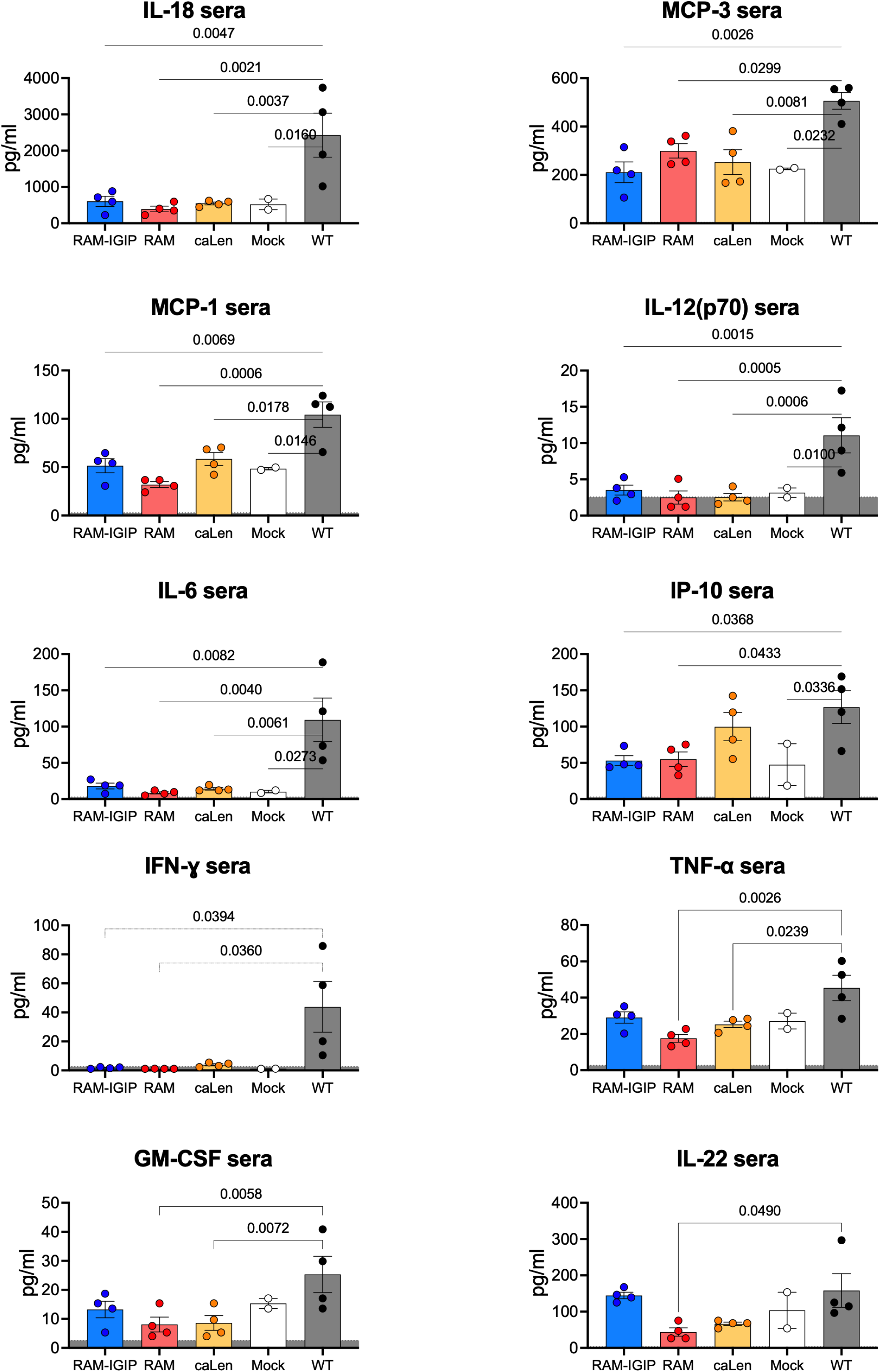
Innate immune responses in serum after virus challenge. Serum cytokine levels were analyzed in samples collected at 5 dpc. Only those with statistically significant differences among groups are shown. Values in y-axis correspond to pg/mL. Statistically significant differences are shown with p values above lines or brackets.

### FLUAV RAM and FLUAV RAM-IGIP MLVs elicit neutralizing and antibody dependent cytotoxic responses stimulated after boost

A subset of previously vaccinated mice (n=4/group, ½ female) received a boost of the same vaccine at 21 dpi. Humoral immune responses were evaluated at 20 dpi (prime) and 14 days after boost (prime-boost, Fig 7). Sera from 4 mice per group/time point were analyzed. After prime immunization, HI titers (Fig 7A) were detected in serum samples from both the RAM-IGIP group (HI=20, 40, 40, 80) and RAM group (HI 20, 40, 40, 40). The caLen group samples had HI titers = 10. HI titers increased significantly after boost: RAM-IGIP with HI=40, 160, 320, 320, RAM with HI=40, 80, 80, 160, and caLen with HI=10, 20, 40, 80. As expected, mock control samples had HI titers <10 at either after prime or boost. Neutralizing antibodies were also screened by a modified microneutralization assays (MN) using a recombinant H1N1 FLUAV virus expressing nanoluciferase (Fig 7B) (12). In this assay, the levels of NLuc activity are inversely proportional to the levels of neutralizing antibodies. After the boost, the reduction in NLuc activity was more evident in the RAM-IGIP and RAM groups compared with the caLen group. The log2 of the inhibitory serum dilution 50 (ISD50) was 9.74, 8.57, and 6.07 for RAM-IGIP, RAM, caLen, respectively. Next, we assessed the levels of antibody-dependent cellular cytotoxicity (ADCC) against the homologous virus in sera samples after boost (Fig 7C). Statistically significant differences were observed between RAM-IGIP and RAM groups compared to the caLen and mock-vaccinated groups.

**Fig 7.**
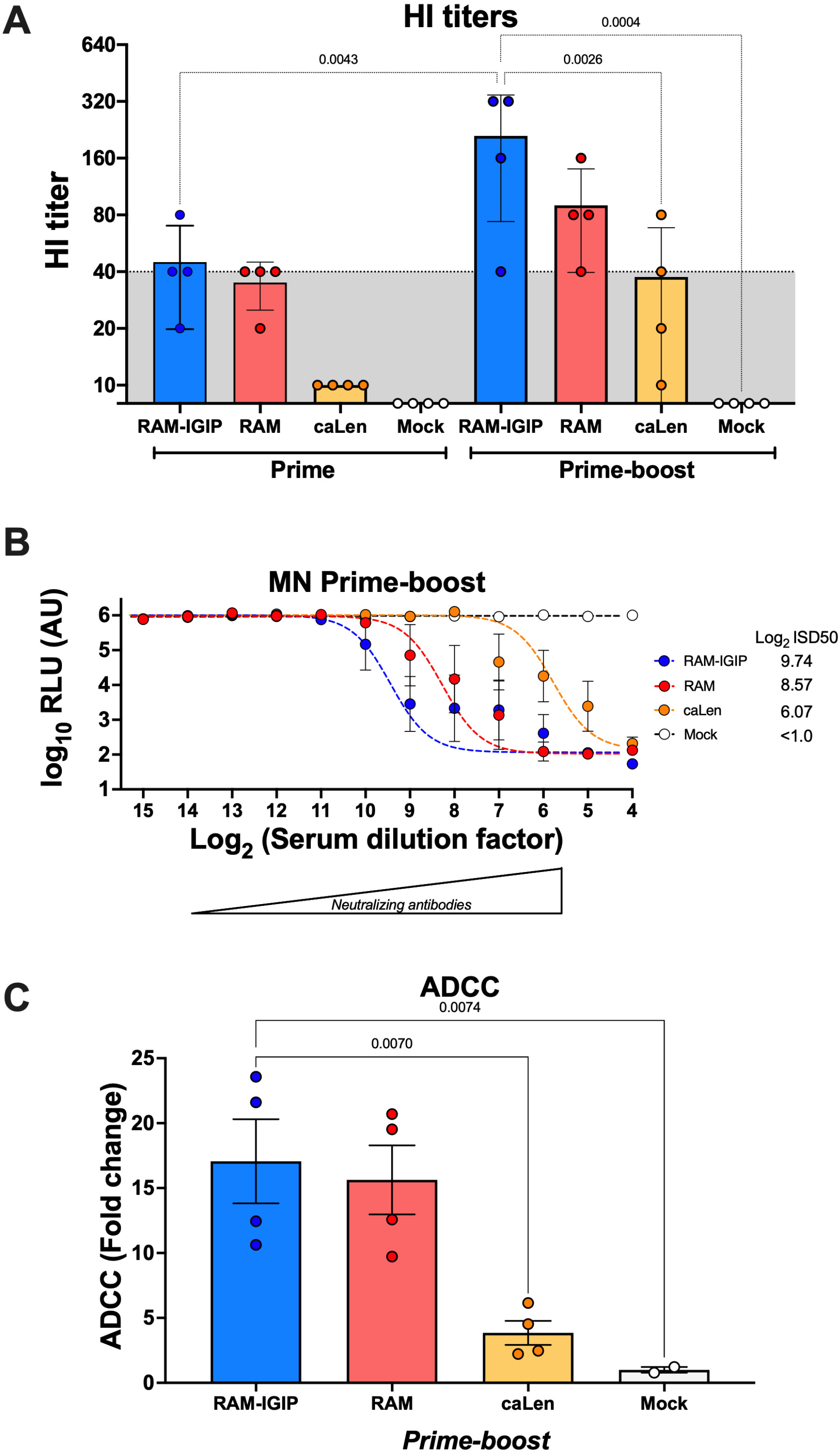
Neutralizing responses post-prime and post-boost. Mice received one or two doses of the same vaccine, 3 weeks apart. **(A)** Hemagglutination inhibition (HI) at 20 dpi and 14 dpb are shown. **(B)** Microneutralization (MN) using a modified Nluc assay (VNluc) is shown on serum samples collected at 14 dpb. **(C)** Antibody-dependent cellular cytotoxicity. Data was normalized to values from mock-infected animals and expressed as fold changes.

### Broadly reactive serum IgG responses associated with RAM-IGIP samples

Total IgG responses against various influenza HA subtypes were analyzed using a protein microarray containing HAs from group 1 (n=71 proteins) and group 2 (n=66 proteins) FLUAV subtypes (as well as from the two major FLUBV lineages as controls, n=22 proteins) (Table S1) (12, 18). Approximately half of the HA proteins on the array are represented as full-length sequences, while the remaining correspond to the HA1 region. Post-prime and post-boost IgG responses against H1 antigens (Fig 8A) were significantly higher in serum samples from the RAM-IGIP and RAM groups compared to the caLen group. Responses against H5 and H9 antigens were lower compared to H1 responses, as reflected in the differences in the MFI signals on the y-axis, but RAM-IGIP-vaccinated mice exhibited elevated anti-H5, and anti-H9 IgG responses compared to other vaccine groups (Fig 8B-C). Overall, these findings demonstrate that the RAM MLVs induce robust IgG responses in serum, with RAM-IGIP providing enhanced cross-reactive responses to other group 1 HA antigens.

**Fig 8.**
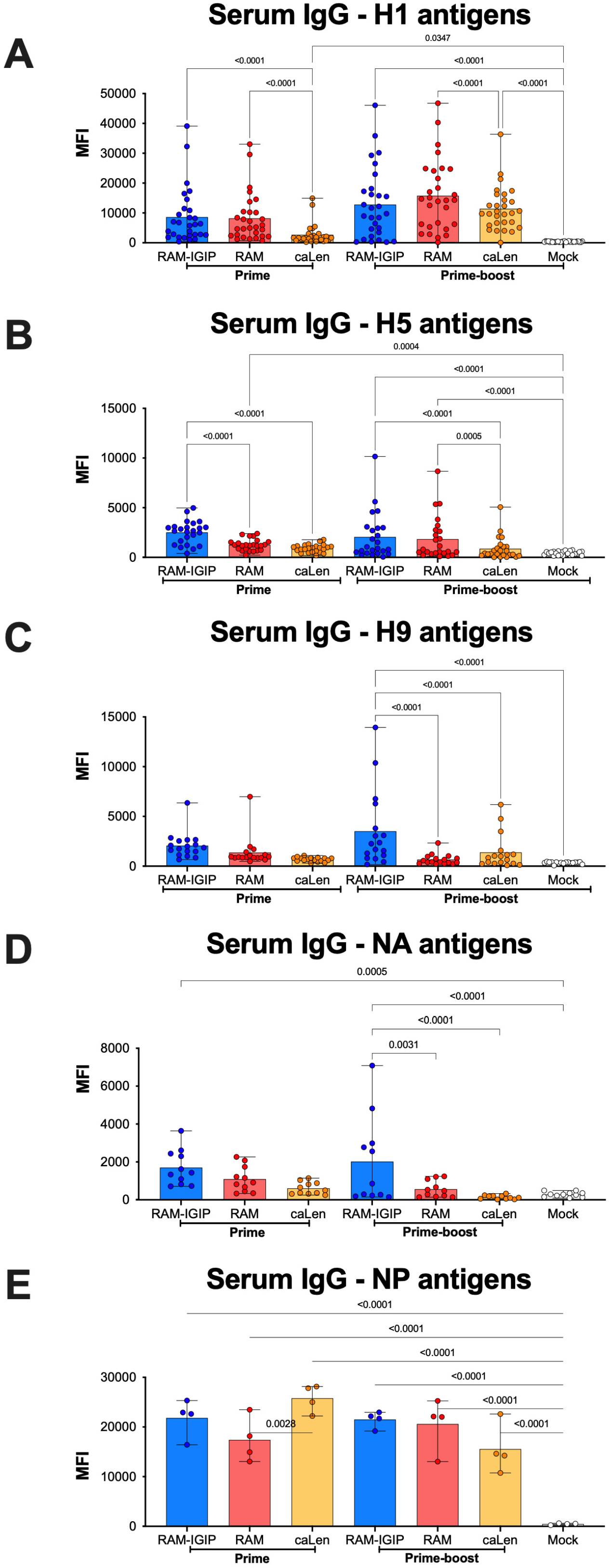
Serum IgG responses against Group 1, NA, and NP antigens. Mice received one or two doses of the same vaccine 3 weeks apart. Detection of IgG from serum was performed against a panel of **(A)** H1. **(B)** H5. **(C)** H9. **(D)** NA and **(E)** NP antigens using an influenza antigen microarray.

Of note, serum IgG responses against Group 2 HA were detected, but the level of detection varied depending on the subtype, vaccine group, and time point of sample collection (Fig S4). Serum IgG against H7 HA increased significantly after boost for the RAM-IGIP and RAM groups but not for the caLen group. The RAM group had the largest anti-H7 effect after boost. In contrast, serum IgG against H3 were more prominent post prime, particularly in the RAM-IGIP group, but all vaccine groups showed significant drops in MFI signal after boost.

The microarray analysis also facilitated the evaluation of immune responses against 12 NA proteins representing the N1, N2, and N9 subtypes and 4 different NP proteins (Figure 8D-E, Table S1). Serum IgG responses against NA antigens (Figure 8D) revealed an elevation in anti-NA IgG antibodies in samples from RAM-IGIP, particularly after the boost, compared to RAM and caLen groups. As anticipated, the most reactive antigens were against the N1 NA proteins on the array, especially from A/California/04/2009 (H1N1) and A/Hubei/1/2011 (H5N1) (data not shown). MFI signals suggest that responses against NP were significantly higher compared to mock control serum samples (Figure 8E). Interestingly, the caLen group produced the strongest anti-NP responses after the prime but the weakest recall responses after the boost. In contrast, anti-NP responses from the RAM-IGIP and RAM groups remained largely unchanged after both prime and boost vaccinations.

### RAM-IGIP associated with enhance mucosal IgG and IgA responses

Mucosal immune responses were investigated using bronchoalveolar lavage fluid (BALF) collected after prime and boost vaccinations (Fig. 9). Mice vaccinated with RAM-IGIP exhibited the highest mucosal anti-H1 HA IgG levels at both prime and boost, indicating a strong immune response against H1N1 influenza virus. Similar patterns were observed for anti-H5 and anti-H9 IgG responses, with significantly higher IgG levels in RAM-IGIP-vaccinated mice compared to other groups, suggesting broader mucosal immunity against H5 and H9 viruses. As expected, responses against H1 were considerably higher than H5 and H9 responses, reflecting the focus of the vaccination strategy.

**Fig 9.**
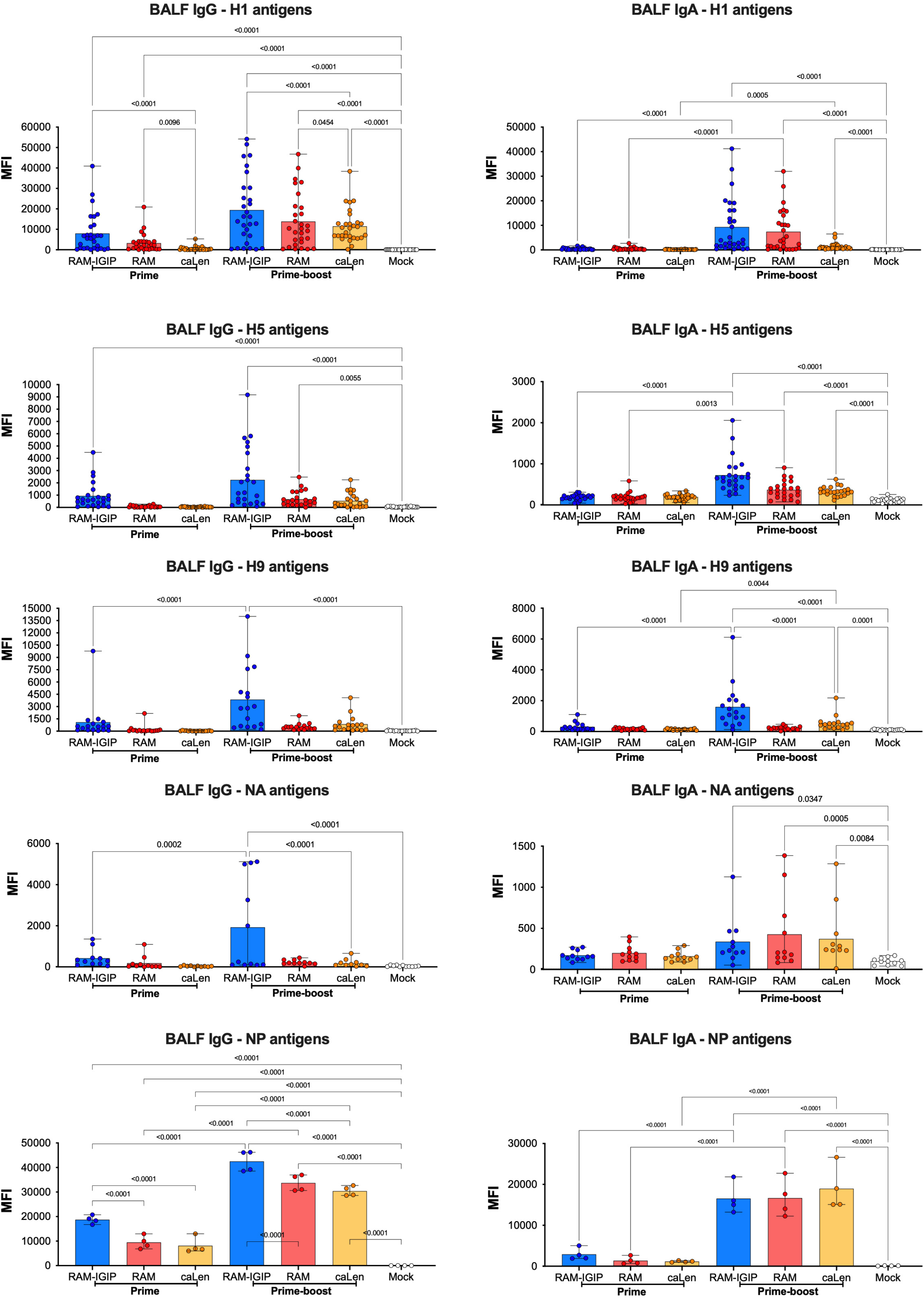
Mucosal IgG and IgA responses against Group 1, NA, and NP antigens. Mice received one or two doses of the same vaccine 3 weeks apart. Detection of mucosal IgG and IgA using BALF samples was performed against a panel of H1, H5, H9, NA and NP antigens using an influenza antigen microarray.

While mucosal IgA levels appeared lower than IgG based on MFI signals, the response pattern against H1, H5, and H9 antigens mirrored that of IgG responses, further supporting mucosal immunity against these viruses. Anti-HA IgA levels in BALF samples were initially low after prime but increased significantly after boost with both RAM-based MLVs, particularly against H1 HA antigens, indicating an effective IgA response elicited by the vaccination regimen.

For group 2 HA antigens, BALF IgG and IgA responses followed a similar pattern as serum IgG responses (Fig S4). Significant increases in anti-H7 responses after boost were detected in RAM-IGIP and RAM samples, while caLen samples showed negligible responses. Against H3, BALF IgG and IgA signals remained moderately above background despite some statistical differences compared to samples from mock-vaccinated mice.

Mucosal IgG anti-NA responses (Fig. 9) were marginal after prime immunization but increased significantly after boost only in the RAM-IGIP group, demonstrating a specific and amplified mucosal immune response against NA in RAM-IGIP-vaccinated mice. In contrast, marginal IgA anti-NA responses were observed, with values close to the background (Fig. 9), suggesting a limited role of mucosal IgA in the immune response against NA in this study.

Mucosal anti-NP IgG and IgA responses (Fig. 9) increased after boost in all vaccine groups. However, BALF IgG anti-NP levels were higher in mice from the RAM-IGIP group after both prime and boost vaccinations, indicating a stronger IgG response against NP in this group. While BALF IgA levels after boost increased considerably across all groups with no significant differences, this mimicked the behavior of anti-NP serum IgG responses, suggesting a broad and consistent IgA response against NP across all vaccination groups.

Overall, the mucosal response analysis demonstrates stronger mucosal IgG and IgA responses in the context of the RAM MLVs, with an enhancement associated with the inclusion of IGIP.

## DISCUSSION

While vaccination remains the primary defense against influenza, overall vaccine efficacy is below 40% (19, 20). LAIVs hold promise for inducing multifaceted and universal cellular and humoral responses. However, they have also been associated with suboptimal efficacy. A notable example is during the 2016-17 and 2017-18 influenza seasons when the Advisory Committee on Immunization Practices (ACIP) recommended that LAIV not be used because of concerns about low effectiveness against the H1N1 viruses circulating in the United States during the 2013-14 and 2015-16 seasons(21). Therefore, there is an urgent need for novel LAIVs for influenza.

We developed a FLUAV genome rearrangement strategy (RAM) that incorporates IGIP into the HA segment of candidate RAM MLVs. We chose IGIP because it is naturally found in mammals and has the potential to boost the immune response against the virus (9, 10). We specifically used the swine IGIP sequence in this study, which is identical to the mature IGIP peptide found in cows, pigs, and ferrets. However, the mature IGIP sequence in mice and humans differs slightly from the swine sequence. The predicted mature IGIP from mouse differs from the swine homolog at position 10 (threonine versus asparagine) and from the human homolog at positions 2 (asparagine versus lysine) and 10 (threonine versus asparagine). The swine and human homologs differ from each other at position 2 (asparagine versus lysine). While the role of these differences is unknown, our previous research showed that they have minimal impact on the immune response in mice (12). We created two versions of the RAM virus: one with IGIP and one without. Both versions remained stable even after repeated passages in tissue culture and eggs.

Safety studies in mice demonstrated that both the RAM and RAM-IGIP viruses were attenuated, like the well-established caLen virus. No observable clinical signs or abnormal behavior were detected in mice infected with either RAM virus, indicating that the RAM rearrangement itself is sufficient to attenuate the virus *in vivo*. These findings align with previous studies exploring various rearrangement strategies, where modifications to PB1, M, NS genes, or incorporation of heterologous sequences successfully attenuated FLUAV viruses in different animal models (5, 22, 23). In the respiratory tract, unlike the caLen group where virus replication was undetectable, both the RAM-IGIP and RAM groups exhibited measurable levels of virus replication. This finding is consistent with previous observations in mice vaccinated with LAIVs derived from the A/Ann Arbor/6/60 donor strain (24). Interestingly, histopathological analysis revealed slightly reduced attenuation in the RAM-IGIP virus compared to other vaccine groups. This was reflected in a trend towards higher levels of pro-inflammatory cytokines in both serum (MCP-1, IL-18, IL-12(p70), TNF-α, GM-CSF, IL-6, MCP-3, and IL-2) and lungs (GRO-α and TNF-α). While these cytokine levels were significantly lower than those observed in control mice infected with WT Ca/04 virus, their stimulation could potentially contribute to the development of protective immune responses against influenza (25–30). This response might be mediated by the recruitment of antigen-presenting cells, natural killer cells, and CD4+ and CD8+ T cells (25–30). The increased pro-inflammatory cytokines observed in RAM-IGIP mice could be attributed to either the activity of IGIP within the respiratory tract or a potential difference in the virus particle/infectious unit ratio compared to other vaccine strains. However, the lower RAM-IGIP replication levels at 5 dpi in various tissues suggest the latter is less likely. Despite extensive efforts, we were unable to detect any IGIP expression from the modified HA protein, possibly due to either alternative translation initiation bypassing the IGIP sequence (222 nt downstream of the first ATG on a second ATG codon immediately downstream of the 2ATav sequence and corresponding to the Gaussia luciferase signal peptide sequence). The observed immune response observed with RAM-IGIP vaccine could be solely attributable to the modifications made to the HA segment itself. Further research beyond the scope of this report employing different IGIP expression detection methods and comparison with alternative irrelevant peptide sequences is needed to definitively determine the cause of the observed immune response.

Both RAM-IGIP and RAM vaccinated groups exhibited complete protection against the challenge, as evidenced by the absence of clinical signs and weight loss (Fig. 4A-D). In contrast, caLen vaccinated mice displayed mild clinical symptoms, including significant body weight loss between 3 and 5 dpc. While live virus was undetectable after challenge in all vaccinated mice, analysis of vRNA loads revealed differences in protection. Overall, vRNA levels were higher in caLen vaccinated mice compared to the RAM-IGIP or RAM groups. The most significant difference was observed in the lungs, where higher FLUAV replication is associated with increased disease severity (31). Furthermore, RAM-IGIP and RAM vaccinated groups showed significantly lower expression of pro-inflammatory cytokines in the lungs at 5 dpc compared to caLen and/or mock-vaccinated-challenged groups (Fig 5). This included cytokines such as IL-1ß, IP-10, MIP-1α, GM-CSF, IL17A, TNF-α, MIP-1 ß, MCP-3, IL-22, and GRO-α. Additionally, the RAM-IGIP group displayed lower expression of MCP-1, IL-6, IL-18, IFN-γ, IL-12(p70), IL-23, IL-10, and IL-2 compared to mock-vaccinated-challenged controls. The RAM and caLen groups showed an intermediate phenotype in this latter set of lung cytokines, with lower levels than mock-vaccinated-challenged controls, although less pronounced in the caLen group. In sera at 5 dpc, all vaccine groups exhibited lower pro-inflammatory cytokine expression compared to mock-vaccinated-challenged controls (Fig 6). However, some exceptions were observed: IP-10 serum levels remained similar between the caLen and mock-vaccinated-challenged groups. TNF-α, GM-CSF, and IL-22 levels were comparable between the RAM-IGIP and mock-vaccinated-challenged groups. The elevated levels of lung and serum IP-10 (as well as other pro-inflammatory cytokines) observed in caLen-vaccinated mice align with previous reports linking active virus replication and disease severity during influenza and SARS-CoV-2 infections (29, 32-34). Interestingly, lower levels of lung IL-23, IL-10, and IL-2, and increased serum levels of TNF-α, GM-CSF, and IL-22 in the RAM-IGIP group are consistent with improved protection against influenza. Overall, these findings suggest that RAM-IGIP and RAM vaccines offer superior protection against influenza compared to the caLen vaccine. This enhanced protection is likely associated with reduced vRNA loads and pro-inflammatory cytokine expression, particularly in the lungs.

RAM-IGIP and RAM vaccines elicited consistently higher levels of antibodies against H1 HA compared to the caLen group, with a trend towards even higher levels in the RAM-IGIP group (Fig. 7-9, S4). The caLen responses align with the low neutralizing antibody responses typically observed in LAIVs approved for human use, where efficacy is measured directly rather than relying on HI titers as correlate of protection (e.g., HI titer ≥ 40) (35). Total IgG analysis via influenza antigen microarray confirmed these findings (Fig 8), revealing higher responses in the RAM-IGIP and RAM groups against H1 antigens. Notably, the RAM-IGIP group displayed superior responses against H5 and H9 antigens, especially after two doses. This suggests that incorporating IGIP into the HA segment modulates the IgG immune response, potentially broadening its target range.

Mucosal immune responses further supported the superiority of the RAM-IGIP vaccine (Fig 9). RAM-IGIP vaccinated mice showed significantly higher IgG and IgA antibody levels against H1, H5, and H9 antigens compared to other groups. Responses against H3 antigens were generally low across all groups. However, RAM-IGIP and RAM vaccinated mice showed a unique response against H7 antigens (Fig S4). Regarding more conserved FLUAV epitopes like NA and NP (Fig. 8 and 9), RAM-IGIP elicited a robust IgG and IgA responses, similar or superior to other vaccine groups. This highlights the potential of RAM-IGIP to induce broader immune protection. This reinforces the potential of IGIP to modulate the immune response in ways yet to be fully understood. While IGIP typically requires CD40L for IgA class switching in bovine B cells, it can also stimulate IgA production in human B cells (9, 11). Interestingly, the observed association between IGIP and IgG production is unexpected, as previous studies have not reported such effects.

Several strategies for developing universal influenza vaccines target conserved epitopes on NA, M2, M1, and NP (36–39). NP, for instance, modulates immune responses by activating CD4+ and CD8+ lymphocytes, potentially offering cross-reactivity against zoonotic influenza strains (40, 41). Studies on LAIVs have shown that different NPs can modulate the immune response differently, leading to protection against heterologous challenges even in the absence of neutralizing antibodies (42). Anti-NA antibodies, although less studied than HA antibodies, have been shown to correlate with protection and reduced viral shedding in humans (43, 44). NA antibodies have also been demonstrated to confer protection and prevent FLUAV transmission in animal models (45). Recent research suggests that N1 antibodies from H1N1-infected individuals can provide heterologous protection against H5N1 (46).

These findings collectively highlight the advantages of a vaccine system that stimulates the production of antibodies and cell-mediated responses against multiple viral antigens, as exemplified by the RAM-IGIP vaccine.

Combining a novel genome rearrangement strategy (RAM) with the immunomodulatory peptide IGIP, we developed a new MLV influenza vaccine. RAM-IGIP showed *in vivo* attenuation and conferred complete protection against a lethal challenge. Additionally, cytokine profiles suggested IGIP enhanced immune responses and overall MLV performance. Future research aims to clarify IGIP’s role in immunogenicity across different vaccine platforms, including mRNA and vector-based vaccines. This work paves the way for developing next-generation influenza vaccines with integrated immunomodulators, contributing to the quest for universal influenza vaccines.

## MATERIALS AND METHODS

### Cells

Madin-Darby canine kidney (MDCK) and human embryonic kidney 293T cells (HEK293T) were a kind gift from Robert Webster (St Jude Children’s Research Hospital, Memphis, TN). Cells were maintained in Dulbecco’s Modified Eagles Medium (DMEM, Sigma-Aldrich, St Louis, MO) containing 10% fetal bovine serum (FBS, Sigma-Aldrich, St Louis, MO), 1% antibiotic/antimycotic (AB, Sigma-Aldrich) and 1% L-Glutamine (Sigma-Aldrich). Cells were cultured at 37°C under 5% CO2. Human airway epithelial (HAE) cells BCi-NS1.1 were obtained from Dr. Matthew S. Walters (Weill Cornell Medicine, NY, USA). Cells were maintained in 1X Basal Media containing PneumaCult-Ex Plus Basal Medium (490mL) (STEMCELL Technologies, Vancouver, Canada), PneumaCult-Ex Plus 50X Supplement (STEMCELL Technologies), 0.1% Hydrocortisone (STEMCELL Technologies), 1% Penicillin-Streptomycin (5000 U/mL; ThermoFisher Scientific, MA, USA), 0.5% Amphotericin B (ThermoFisher Scientific), and 0.5% Gentamycin (50 mg/mL; Sigma-Aldrich). Cells were cultured at 37°C under 5% CO2. Differentiation of the BCi-NS1.1 was performed in 12.5 mm trans-well plates with 0.4 um pore polyester membrane inserts (Corning Inc., NY, USA). Before plating, trans-well membranes were coated with human type IV collagen (Sigma-Aldrich, St. Louis, MO, USA) and then rinsed with 1X phosphate-buffered saline (PBS; ThermoFisher Scientific). Once the membrane was dry, 300,000 BCi-NS1.1 cells/well were plated with 1X Basal Media and cultured at 37°C under 8% CO2. After reaching 100% confluency, the cells were changed to ALI conditions by removing the apical media and then changing the basal media for 1X ALI media. ALI media contained PneumaCult ALI Base Medium (450mL) (STEMCELL Technologies), 10X PneumaCult ALI Supplement (50mL) (STEMCELL Technologies), 1% Penicillin-Streptomycin (5000 U/mL; ThermoFisher Scientific, MA, USA), 0.5% Amphotericin B (ThermoFisher Scientific, MA, USA), 0.5% Gentamycin (50 mg/mL; Sigma-Aldrich), 1% PneumaCult ALI Maintenance Supplement (STEMCELL Technologies), 0.2% Heparin solution (STEMCELL Technologies, Vancouver, Canada), and 0.5% Hydrocortisone Stock solution (STEMCELL Technologies). Cells were cultured in ALI conditions at 37°C under 8% CO2 for 5 days and after at 37°C under 5% CO2 until they reached 28 days in ALI conditions.

### Generation of plasmid constructs IGIP-HA, PB1-M2, and M_esc_

The generation of IGIP-H1 has been previously described (12, 47). Briefly, DNA fragments with the sequence corresponding to the 5’ untranslated region (UTR) and signal peptide sequence of H1 HA (A/California/04/09 (Ca/04) (H1N1)), followed by a G4S linker, furin cleavage site, the Thosea assigna virus (TAV) 2A protease and the mature IGIP was generated with a cloning spacer downstream and acquired from Genscript (Piscataway, NJ). The fragment was digested with AarI (Thermo Scientific) and cloned into the reverse genetics plasmid vector pDP2002. For the generation of pDP2002-PB1-M2, DNA fragments with the sequence corresponding to the 5’ untranslated region (UTR), the PB1 open reading frame (ORF) from A/California/04/09 (H1N1) (Ca/04), followed by a G4S linker, the Thosea assigna virus (TAV) 2A protease, M2 ORF of Ca/04 and 83 nucleotides of the 3’ end of the PB1 gene (40 nucleotides encoding the C-terminus of PB1 ORF and 43 from the 3’UTR region) was synthesized by Genscript (Piscataway, NJ). The fragments were digested with BsmBI, purified, and cloned into pDP2002, as previously described (48). The plasmid pDP2002-M_esc_ with early stop codons in the M2 ORF was generated by PCR introducing the sequence 5’-taatgataatag-3’ between positions 786-797 of the M gene (amino acid positions 25-28 of the M2 ORF). All plasmid sequences were confirmed by Sanger sequencing (Psomagen, Rockville, MD).

### Generation of RAM viruses by reverse genetics

The Ca/04-derived pDP2002-IGIP-H1 or the pDPHA-H1 (Ca/04) WT plasmids were transfected together with pDP2002-NA (Ca/04), pDP2002-PB1-M2, pDP2002-Mesc, and the remaining plasmids (PB2, PA, NP, and NS) derived from the WT OH/04 backbone previously described (15, 49). The caLen virus was previously described and obtained using pDPHA-H1 (Ca/04) WT and pDP2002-NA (Ca/04) and the internal gene segments corresponding to the cold-adapted Leningrad backbone (caLen), kindly provided by Irina Isakova-Sivak and Larisa Rudenko, Department of Virology, Institute of Experimental Medicine, St Petersburg, Russia (12, 17). Attempts to rescue an IGIP-H1 construct in the background of caLen led to a virus with low yields (12). Reverse genetics was performed in co-cultured HEK293T and MDCK cells as previously described (50). Viral stocks were generated in 10-day-old specific pathogen-free eggs. Allantoic fluids were harvested 48 h post-infection (hpi), cleared by centrifugation, aliquoted, and stored at −80°C. Viruses were titrated by tissue culture infectious dose 50 (TCID50), and virus titers were established by the Reed and Muench method (51). Virus sequences were confirmed by next-generation sequencing and Sanger sequencing, as previously described (6, 52)

### In vitro growth kinetics

MDCK (6-well plate) or differentiated HAE (12-well plate) cells were inoculated with the different viruses at an MOI of 0.01. Cells were incubated with the inoculum for 15 min at 4°C and then 45 min at 35°C. Subsequently, the inoculum was removed, and MDCK and HAE cells were washed with PBS. MDCK cells were cultured in Opti-MEM media containing 1% AB while HAE cells were cultured in ALI media. Cells were incubated at 35°C under 5% CO2, and supernatant samples were collected at 0, 12, 24, 48, and 72 h post-inoculation and stored at −80°C until further processing. Supernatants from HAE cells were collected from the apical side by adding/removing 200 µL of Opti-MEM media containing 1% AB.

### Mouse studies

Animal studies were approved by the Institutional Animal Care and Use Committee at the University of Georgia (AUP: A2019 03-032-Y3-A16) and performed under animal biosafety level 2 conditions. Animal studies and procedures were performed according to the Institutional Animal Care and Use Committee Guidebook of the Office of Laboratory Animal Welfare and PHS Policy on Humane Care and Use of Laboratory Animals. Animal studies were carried out in compliance with the ARRIVE guidelines (https://arriveguidelines.org). Female and male Balb/c mice (5 to 6 weeks old) were purchased from Jackson Laboratories (Bar Harbor, ME). Mice were randomly distributed into 5 groups: WT Ca/04, RAM-IGIP, RAM, caLen, and mock vaccinated. Animals were anesthetized with isoflurane and inoculated intranasally (I.N.) with 50 µL of either phosphate buffer saline (PBS; mock) or 1×10^5^ TCID50/mouse of the different viruses. At 5- and 20-dpi, a subset of mice (n=4/group/time point) were anesthetized, bled, and humanely euthanized to collect different fluids and tissues. At 21 dpi, a subset of mice (n=12/group) were challenged with 1×10^6^ TCID50/mouse (∼1,000 mice lethal dose 50) of WT Ca/04 (14, 53). At 5 days post-challenge (5 dpc), a subset of mice (n=4/group, n=2/mock unvaccinated non-challenged control) were anesthetized, bled, and humanely euthanized to collect sera and different tissues. In addition to the prime-challenge regime, 4 mice per group were boosted at 21 dpi with the same vaccine and dose used at prime. Mice were monitored along the entire course of the experiments for clinical signs at least once daily. Mice that lost ≥ 25% of their initial body weight (and/or a score of 3 on a 3-point disease severity scale) were humanely euthanized. To obtain serum samples before euthanasia, mice were bled from the submandibular vein as previously described (54). Thirty-five days after the beginning of the experiment, all remaining mice were anesthetized, bled, and humanely euthanized, and the experiment was terminated.

### Viral RNA amplification and quantification

Virus RNA from *in vitro* or *in vivo* experiments was isolated using the QIAamp® Vira RNA Mini Kit (Qiagen, Germantown, MD) or the MagMAX-96 AI/ND viral RNA isolation kit (Thermo Fisher Scientific) following the manufacturer’s protocol. RT-PCR assays were performed using the SuperScript III One-Step RT-PCR System with Platinum Taq DNA Polymerase (ThermoFisher Scientific) following manufacturer’s conditions. Amplicons were resolved in a 1% agarose gel and visualized using the ChemiDoc MP Imaging System (Biorad, Hercules, CA). When specified, virus titers were determined using a real-time reverse transcriptase PCR (RT-qPCR) assay based on the influenza A matrix gene. The RT-qPCR was performed in a QuantStudio 3 (Applied Biosystems, Foster City, CA) using qScript XLT One-Step RT-qPCR ToughMix®, QuantaBio (ThermoFisher). A standard curve was generated using 10-fold serial dilutions from a virus stock of known titer to correlate quantitative PCR (qPCR) crossing-point (Cp) values with virus titers, as previously described (12, 55). Virus titers were expressed as log10 TCID50 equivalents per mL of tissue homogenate supernatant as noted.

### Hemagglutination inhibition assays

Serum samples were collected at 20 dpi and 14 dpb to screen for the presence of neutralizing antibodies by hemagglutination inhibition (HAI) assays as previously described (53). Briefly, the sera were treated with receptor-destroying enzyme (Denka Seiken, VWR), incubated overnight at 37°C, and then inactivated at 56°C for 30 min. After inactivation, the sera were diluted 1:10 with PBS, serially diluted 2-fold, and mixed with 4 hemagglutination units (HAU) of virus in a 96-well plate. The virus-sera mixture was incubated for 15 min at room temperature, and the HI activity was determined after 45 min of incubation with 0.5% of turkey red blood cells (RBC). HI titers below ≤10 was arbitrarily assigned a value of 8.

### Virus neutralization assays

Microneutralization assays were performed as previously described with a modification utilizing a nanoluciferase detection method (56, 57). Briefly, the recombinant Ca/04 (H1N1) virus carrying Nano luciferase (NLuc) gene downstream the PB1 segment was used at 100 TCID50 per well in a 96-well plate and incubated with 1/10 serial dilutions of serum samples collected and treated as described above. The serum-virus mixture was incubated for 1 h at 35°C, overlayed for 15 min at 4 °C, and then 45 min at 35°C on MDCK cells seeded in a 96-well plate the day before. The serum-virus mixture was subsequently removed, and 200 µL of Opti-MEM-AB + TPCK-trypsin was added. The cells were incubated at 35°C under 5% CO2 for 48 h. The virus neutralization titers were determined by measuring the NLuc activity using the Nano-Glo Luciferase Assay System (Promega, Madison, WI) following the manufacturer’s conditions. NLuc activity was quantified in a Victor X3 multilabel plate reader (PerkinElmer, Waltham, MA).

### Antibody-dependent cell-mediated cytotoxicity (ADCC)

The ADCC activity from sera samples collected at 20 dpi and 14 days post-boost (dpb) was measured using the mFcyRIV ADCC Reporter Bioassay (Promega). MDCK cells were seeded in a 96-well plate at 20,000 cells/well the day before the assay. The next day, media was removed, and cells were infected with Ca/04 at a MOI=5 and incubated for 24 h. After the incubation, cell culture media was removed, and 50 μl of serum diluted in ADCC assay buffer (1/10 dilutions) pre-heated at 56°C for 30 min was added to the cells. Next, 25μl of effector cells was added to each well following the manufacturer’s conditions. Plates were covered and incubated for 6 h at 35°C under 5% CO2. After the incubation, the levels of luciferase activity were measured using 70 μl of the Bio-Glo reagent following the manufacturer’s conditions. Luciferase activity was quantified in a Victor X3 multilabel plate reader (PerkinElmer).

### Tissue homogenates preparation

Tissues (Nasal turbinates, trachea, lungs, and brain) homogenates collected from mice at 5 dpi and 5 dpc were generated using the Tissue Lyzer II (Qiagen). Briefly, 1 mL of PBS-AB was added to each tissue with Tungsten carbide 3mm beads (Qiagen). Samples were homogenized for 15 min and then centrifuged at 15,000 g for 10 min at 4°C. Supernatants were collected, aliquoted, and stored at −80 until further analysis. Samples were titrated by TCID50, and virus titers were established by the Reed and Muench method (51).

### Luminex assays

Analysis of cytokines and chemokines from sera and lungs homogenates collected 5 dpi and 5 dpc were conducted using the ProcartaPlex TM Mouse Cytokine & Chemokine Convenience Panel 1 26-plex (Thermo Fisher Scientific) following the manufacturer’s specifications. The capture beads mix was briefly vortexed, and 50 μl was added to each well in a 96-well plate. After washing, serum (1/2 dilution in UAB buffer) and lungs homogenates (undiluted) were added to the plate, shaken at 600rpm for 30 min at room temperature, and then transferred at 4°C overnight. Next, the plate was placed on the hand-held magnetic plate, washed 3 times, and 25 μl of detection antibody was added to each well. The plate was sealed and incubated at 600 rpm for 30 min at room temperature. After the incubation, the plate was washed 3 times and 50 μl of the Streptavidin-PE antibody was added to each well, and the plate was incubated at 600 rpm for 30 min at room temperature. Finally, 120 µL of the reading buffer was added per well; the plate was set at 600rpm for 5 min at room temperature. The expression of each cytokine was captured using a Luminex MAGPIX® instrument.

### Influenza antigen microarray

The influenza antigen microarray was performed as previously described [23]. Serum and BALF samples were diluted 1:100 in protein array blocking buffer (GVS, Sanford, ME, USA) supplemented with E. coli lysate (GenScript, Piscataway, NJ, USA) to a final concentration of 10 mg/mL and preincubated at room temperature (RT) for 30 min. Concurrently, arrays were rehydrated in blocking buffer (without lysate) for 30 min. The blocking buffer was removed, and arrays were probed with preincubated serum samples using sealed chambers to prevent cross-contamination of samples between the pads. Arrays were incubated overnight at 4°C with gentle agitation. They were then washed at RT three times with Tris-buffered saline (TBS) containing 0.05% Tween 20 (T-TBS), biotin-conjugated goat anti-mouse IgA, and Biotin –conjugated anti-mouse IgG (Jackson Immuno Research Laboratories, Inc., West Grove, PA, USA) were diluted 1:400 in blocking buffer and applied to separate arrays for 1 h, RT with gentle agitation. Arrays were washed three times with T-TBS, followed by incubation with streptavidin-conjugated Qdot655 (Thermo Fisher Scientific, Waltham, MA, USA) diluted 1:200 in blocking buffer for 1 h, RT. Arrays were washed three times with T-TBS and once with water. Arrays were air dried by centrifugation at 500 g for 5 min. Images were acquired using the ArrayCAM imaging system from Grace Bio-Labs (Bend, OR). Spot and background intensities were measured using an annotated grid (.gal) file. Mean fluorescence across antigens grouped by isotypes was used for subsequent analysis. The different antigens were acquired from Sino biological (Wayne, PA).

### Graphs/Statistical analyses

All data analyses and graphs were performed using GraphPad Prism software version 10 (GraphPad Software Inc., San Diego, CA). One-way and/or 2-way ANOVA with multiple comparisons were performed. A P value below 0.05 was considered significant.

## ACKNOWLEDGMENTS

We thank Kristine R. Wilcox and the Life Sciences Animal Facility personnel, University of Georgia. We also thank James Barber and the CVM Cytometry Core Facility personnel, College of Veterinary Medicine, University of Georgia. We are also grateful to Lisa Stabler and the Histology laboratory personnel at the Poultry Diagnostic Research Center, College of Veterinary Medicine, University of Georgia. We thank Irina Isakova-Sivak and Larisa Rudenko, Department of Virology, Institute of Experimental Medicine, St Petersburg, Russia for contributing with reverse genetics plasmids for the caLen vaccine strain.

## DATA AVAILABILITY STATEMENT

Microarray data is deposited in NCBI’s Gene Expression Omnibus(58) and are accessible through GEO Series accession number GSE252919.

## AUTHORS CONTRIBUTION

DRP designed the rearrangement and IGIP strategies. CJC and DPR designed the experiments. CJC performed molecular cloning and rescue of the viruses by reverse genetics. CJC, LCG, and BS performed *in vitro* growth kinetics and stability assays. GG performed viral sequencing. CJC, LCG, BS, BC, FCF, and SC performed *in vivo* studies and sample collection. CJC, LCG, BS TDM, and MC performed *in vivo* data processing. DSR edited the manuscript. SC performed histopathological and immunohistochemistry analyses and edited the manuscript. AJ and HD performed Influenza antigen microarray. CJC and DRP wrote the manuscript. All authors approved the final version of the manuscript.

## Supplementary Materials and Methods

### Polymerase assay

The polymerase activity of the different complexes was determined as previously described (15). Briefly, 300,000 293T cells per well were seeded in 24-well plates in DMEM supplemented with 10% FBS, 1% AB, and 1% L-Glutamine the day before the assay. The following day, cells were transfected with 1 µg each of the pDP2002 plasmids encoding the polymerase complex of RAM (PB2 OH/04, PB1-M2, PA OH/04 and NP OH/04), att (PB2-ts. PB1-att, PA OH/04 and NP OH/04), WT (PB2 OH/04, PB1-M2, PA OH/04 and NP OH/04) or negative control (PB2 OH/04, PA OH/04 and NP OH/04), by using the TransIT-LT1 (Mirus, Madison, WI) reagent according to the recommendations of the manufacturer. In addition, cells were transfected with 1 µg of pNS-GLuc, which was used as a readout of the polymerase activity, and 1 µg of the pCMV/SEAP plasmid, which encodes a secreted alkaline phosphatase (SEAP) gene (total amount of DNA per well = 6 µg). At the indicated time points, 100 µL of supernatants from transfected cells were harvested and assayed for both luciferase and secreted alkaline phosphatase activities by using the BioLux Gaussia luciferase assay kit (New England Biolabs, Ipswich, MA) and the Phospha-Light secreted alkaline phosphatase reporter gene assay system (Applied Biosystems, Foster City, CA) according to the manufacturer’s protocols. Relative polymerase activity was calculated as the ratio of luciferase luminescence to SEAP. Luciferase activity was quantified in a Victor X3 multilabel plate reader (PerkinElmer, Waltham, MA).

### Histopathology examination and virus antigen examination in tissues

Selected tissues were collected from each mouse in each group at 5 dpi and 5 dpc for histopathological examination. Tissues were fixed in 10% neutral-buffered formalin (NBF) for at least 72 h, paraffin-embedded, and processed for routine histopathology with hematoxylin and eosin staining (H&E). A board-certified pathologist, blinded to the study subjectively scored tissues based on % of the total parenchyma affected by lesions and inflammation as none (0%), mild; <15% (1), mild to moderate; 16-30% (2), moderate; 31-50% (3), moderate to severe 51-75% (4) and severe; ≥75% (5). Features considered for the scoring were the following: presence and extent of necrosis, inflammation, epithelial cell hypertrophy/hyperplasia, endothelialitis, mesothelial hyperplasia, presence, and amount of intraluminal catarrhal and suppurative exudate, hemorrhage, edema, fibrin, hyaline membrane formation, and vascular thrombi. For immunohistochemistry (IHC), an antibody targeting the FLUAV nucleoprotein (NP) (ThermoFisher Scientific, Waltham, MA; dilution 1/500) was used on selected tissues. The intensity and distribution of the Fast red chromogen staining was used to estimate the amount of viral NP antigens which was subjectively scored by a pathologist using a scale from none (0) to large amount (5), being the large amount the highest level of positivity.

## Supplementary Figures

**Supplementary Fig 1. Polymerase activity at various temperatures** Polymerase activity assays were conducted with the RAM complex (PB2 WT, PB1-M2, PA WT, and NP WT), att (PB2 ts, PB1 att, PA WT, and NP WT) or a WT complex (PB2 WT, PB1 WT, PA WT, and NP WT). A FLUAV-like segment encoding the (-) RNA of a Gaussia luciferase (GLuc) flanked by the untranslated regions of the NS gene was used as a readout.

**Supplementary Fig 2. Histopathologic findings and virus antigen in trachea and lungs of mice inoculated with MLVs compared to WT Ca/04 and caLen controls at 5 dpi and 5 dpc. (A)** H&E and IHC scores per animal at 5 dpi. **(B)** Representative pictures of the average H&E and IHC scores per group at 5 dpi. **H&E Trachea**: Mice inoculated with WT Ca/04 are the only group with severe suppurative inflammation and accumulation of abundant exudate in the lumen of the tracheas (black arrowhead). RAM-IGIP inoculated mice show mild lymphoplasmacytic infiltration in the tracheal mucosal lamina propria (open triangle). RAM inoculated mice show mild lymphocytic tracheitis (open triangle) at the bifurcation of the trachea with minimal intraluminal catarrhal exudate formation (black arrowhead). The caLen group shows minimal alterations in the trachea, comparable to the Mock controls. **H&E Lungs**: WT Ca/04 inoculated mice show severe suppurative bronchopneumonia with an accumulation of abundant exudate within the bronchi (black arrowhead). Small, sparse clusters of lymphocytes are also infiltrating peribronchial interstitium (open triangle). Mice within the RAM-IGIP group have small foci of atelectasis with minimal pneumocyte type I necrosis and sparse lymphohistiocytic infiltration (asterisk). RAM-inoculated mice have mild-moderate lymphoplasmacytic bronchitis in the lungs (open triangle) with minimal intraluminal catarrhal exudate formation (black arrowhead). Both the caLen group and the mock comparison do not present pulmonary lesions at 5 dpi. IHC Trachea and Lungs: The presence of virus antigen (red staining) is shown by black arrows. The WT Ca/04 inoculated mice present large amounts of virus antigen within intraluminal tracheal and pulmonary bronchial exudate, sparse tracheal epithelial cell nuclei, and pneumocytes type I nuclei. Small to moderate amount of virus antigen staining was observed in the trachea and lungs of mice in the RAM-IGIP group, consistent with viral loads observed. The amount of virus antigen in the RAM group is like the RAM-IGIP group, with slightly less staining in pulmonary samples. The tracheas and lungs of the caLen and mock inoculated group show no evidence of antigen staining. **(C)** H&E and IHC scores per animal at 5 dpc. **(D)** Representative pictures of the average H&E and IHC scores per group at 5dpc. **H&E Trachea**: Moderate suppurative tracheitis with intraluminal tracheal accumulation of exudate (black arrowhead) and minimal lymphocytic inflammation (open triangle) is observed in WT Ca/04 group. The trachea of mice from the RAM-IGIP and RAM groups showed small to moderate numbers of lymphocytes and plasma cells in the tracheal mucosal lamina propria (open triangles). The trachea of the caLen shows lesions similar to the WT Ca/04 of suppurative tracheitis with moderate necrosis of the airway epithelium (black arrowhead). In this group, moderate lymphoplasmacytic infiltrates are expanding the mucosal lamina propria and extending to the overlying respiratory epithelium (open triangles). **H&E Lung**: In the lungs of WT Ca/04 mice, areas of atelectasis, pneumocyte type I necrosis, mixed inflammation of the alveoli (asterisk), and sparse mild infiltration of lymphocytes (open triangles) are observed, in addition to intrabronchial accumulation of suppurative exudate (black arrowhead). The lungs of mice from the RAM-IGIP and RAM groups show small to moderate numbers of lymphocytes and plasma cells forming dense aggregates and lymphoid nodules in the peribronchial pulmonary interstitium (open triangles). The lungs of the caLen show lesions similar to the WT Ca/04 with suppurative bronchopneumonia with necrosis of the airway epithelium (black arrowheads). In the caLen, moderate lymphoplasmacytic infiltrates (open triangles) are more prominent than in the WT Ca/04. IHC Trachea and Lungs: The presence of virus antigen (red) is shown by arrows. The WT Ca/04 inoculated mice present large amounts of virus antigen within intraluminal tracheal and bronchial exudate, sparse tracheal epithelial cell nuclei, and pneumocytes type I nuclei. The caLen group presents small to moderate amount of virus antigen, mostly within tracheal and bronchial intraluminal exudate, and more rarely in the nuclei of the tracheal, bronchial epithelium, and pneumocyte type I (arrows). No virus antigens are detected in the trachea and lungs of the RAM-IGIP and RAM vaccinated groups.

**Supplementary Fig 3. Additional lung cytokines with different expression profiles after challenge depending on vaccine group** Lung cytokines were detected using a 26-plex Luminex assay in samples collected at 5 dpc. Only those with statistically significant differences among groups are shown. Values in y-axis correspond to pg/mL.

**Supplementary Fig 4. Serum IgG and mucosal IgG/IgA profiles against Group 2 HAs** Mice received one or two doses of the same vaccine 3 weeks apart. Detection of serum IgG and mucosal IgG and IgA was performed against a panel of H3 and H7 antigens using an influenza antigen microarray.

